# Metagenomic characterization of the effect of feed additives on the gut microbiome and antibiotic resistome of feedlot cattle

**DOI:** 10.1101/153536

**Authors:** Milton Thomas, Megan Webb, Sudeep Ghimire, Amanda Blair, Kenneth Olson, Gavin John Fenske, Alex Thomas Fonder, Jane Christopher-Hennings, Derek Brake, Joy Scaria

**Affiliations:** Department of Veterinary and Biomedical Sciences, South Dakota State University, Brookings, South Dakota, USA.; South Dakota Center for Biologics Research and Commercialization, South Dakota, USA.; Department of Animal Science, South Dakota State University, Brookings, South Dakota, USA.

## Abstract

In North America, antibiotic feed additives such as monensin and tylosin are added to the finishing diets of feedlot cattle to counter the ill-effects of feeding diets with rapidly digestible carbohydrates. While these feed additives have been proven to improve feed efficiency, and reduce liver abscess incidence, how these products impact the gastrointestinal microbiota is not completely understood. Furthermore, there are concerns that antibiotic feed additives may expand the antibiotic resistome of feedlot cattle by enriching antimicrobial resistance genes in pathogenic and nonpathogenic bacteria in the gut microbiota. In this study, we analyzed the impact of providing antibiotic feed additives to feedlot cattle using metagenome sequencing of treated and untreated animals. Our results indicate that use of antibiotic feed additives does not produce discernable changes at the phylum level however treated cattle had reduced the abundance of gram-positive bacteria at the genus level. The abundance of Ruminococcus, Erysipelotrichaceae and Lachanospira in the gut of treated steers was reduced. This may impact the ability of these animals to exclude pathogens from the gut. However, our results did not show any correlation between the presence of antimicrobial resistance genes in the gut microbiota and the administration of antibiotic feed additives.

## Introduction

Commensal microbes inhabiting the mammalian gut are associated with immune development ^1^, nutrient supply ^2^, prevention of pathogenic diseases ^3^, and overall health of the host. The nutritional dependency on microbial population is particularly important in the case of ruminants as the end products of microbial fermentation in the rumen provide most of the energy required by the animal. Alternatively, dietary practices influence the structure of the microbial community ^4,5^. Cattle in feedlots are generally fed a finishing ration that contains up to 90% rapidly digestible carbohydrates. This elevates incidences of acidosis ^6^ and liver abscesses ^7^, which have a detrimental effect on animal health and beef production. In North America, antibiotic feed additives such as monensin and tylosin are added to the finishing diets to counter the ill-effects of feeding a carbohydrate-rich diet ^8–11^. Monensin is a polyether antibiotic isolated from *Streptomyces cinnamonensis*^12^. Tylosin is a macrolide antibiotic and bacteriostatic feed additive^13^. While these additives are proven to improve feed efficiency, reduce liver abscess incidence, and control coccidiosis, the understanding of how these additives influence the microbial dynamics of the rumen is incomplete. Use of antibiotic feed additives in livestock production is a controversial topic due to the concerns that use of antibiotic feed additives could expand the antibiotic resistome in the animal’s gastrointestinal tract (GIT). The antibiotic resistome is the collection of antibiotic resistance genes and their precursors in both pathogenic and non-pathogenic bacteria in a given environment^14^.

The alterations to the gut microbes in ruminants induced by energy-rich diets and antibiotic feed additives have been studied previously ^6,15–19^. However, the methodologies used in these studies were anaerobic culture, terminal restriction fragment length polymorphism (TRFLP), and 16S rRNA-based polymerase chain reaction (PCR). These techniques lack the ability to analyze whole microbial populations and could only provide information that is limited to few marker organisms. With the recent advances in next-generation sequencing techniques, 16S rRNA gene amplicon sequencing is used to map out the microbial communities in the ruminant gut ^20–25^. However, classifying the microbial community based on a single gene could be challenging and often the resolution is compromised especially when the species are closely related. This method is also inherently limited because of the bias induced by PCR while amplifying the gene. Furthermore, the predicted functional characteristics of the microbial communities using 16S rRNA gene amplicon sequencing might have low accuracy. Shotgun metagenomics sequencing, albeit expensive, is a reliable alternative tool that could provide a comprehensive and high-resolution analysis of the microbiome and resistome.

This study provides a snapshot of the structure and functional characteristics of the GIT microbiome and resistome of steers that are adapted to a rapidly fermentable carbohydrate-rich diet and were finished with or without antibiotic feed additives. To our knowledge, this is the first report that uses shotgun metagenomic sequencing for studying the microbiome and resistome alterations in the GIT of steers. Importantly, the findings of this study are relevant for beef cattle operations and public health since the management and diet of the steers in this experiment resemble the practices followed in the cattle feeding industry in North America.

## Results

In this study, shotgun metagenomics sequencing of the GIT microbiome of antibiotic feed additive treated and untreated feedlot cattle was utilized to determine the influence of these feed technologies on the microbiome and antibiotic resistome. The beneficial effect of feeding antibiotics on improving the health and productivity of feedlot cattle on high-grain diets has been well established. However, it is also a controversial practice that could lead to the emergence of antibiotic-resistant bacterial strains. With the advancement in next generation sequencing technologies, the microbial diversity and antibiotic resistome in the GIT of livestock animals could be studied in greater depth. Previous research conducted on the bovine microbiome made use of a 16S sequencing approach primarily to study the bacterial communities in the GIT. Though it was possible to deduct these results at the operational taxonomic units (OTU), shotgun metagenomics provides better resolution of the taxonomical data to the genera level and even to species level. However, the disadvantage of next generation based metagenome sequencing is that platforms such as Hiseq would have to be used to account for the presence of host genomic DNA contamination. To circumvent this problem, a microbial DNA enrichment method ^26^ was used to deplete the host genomic DNA. Using this method, 49 ±12 %, 53 ±12 %, and 15 ± 4 % of total DNA could be recovered from the rumen, colon, and cecum samples respectively. This depletion process allowed for metagenomics sequencing using 2x 250 paired chemistry in Miseq platform, which provides a 100 base longer read than the maximum length available in the HiSeq platform.

### Diversity and richness estimates

The diversity and richness of the microbial population in the rumen, colon, and cecum samples was obtained from five control steers (NA; not provided any antibiotics or growth promoting treatments) and five steers that were provided antibiotic feed additives and growth promoting implants and beta-agonists (AB). Biodiversity of the human gut is reduced by both short-term and longterm use of antibiotics ^27,28^. The AB steers were fed antibiotic feed additives, monensin and tylosin, which are known to cause changes in the bacterial population in the animal gut ^29,30^. In the present study, we used shotgun metagenomics to study the alterations in the microbial community structure in the GIT of feedlot steers that were fed monensin and tylosin.

### α diversity

The individual sample richness and diversity of rumen, cecum, and colon samples were measured using Chao1, ShannonH, and Inverse Simpson indices by bootstrapping random sequences from the sequences in each sample for 50 times (Table 1). The higher values for ShannonH and inverse Simpson indices indicate that rumen samples from NA steers had a more diversified microbial community. Also, rumen samples from NA steers exhibited higher species richness, as measured by the Chao1 index, than those from AB steers. This suggested that antibiotic treatment might have reduced both community diversity and richness of the microbial population in the rumen. Interestingly, species richness was higher in the colon of AB steers. However, there was no difference observed for richness of the microbial population in the cecum between the treatment groups. Communities had similar diversity in the cecum and colon for both treatment groups.

**Table 1.**
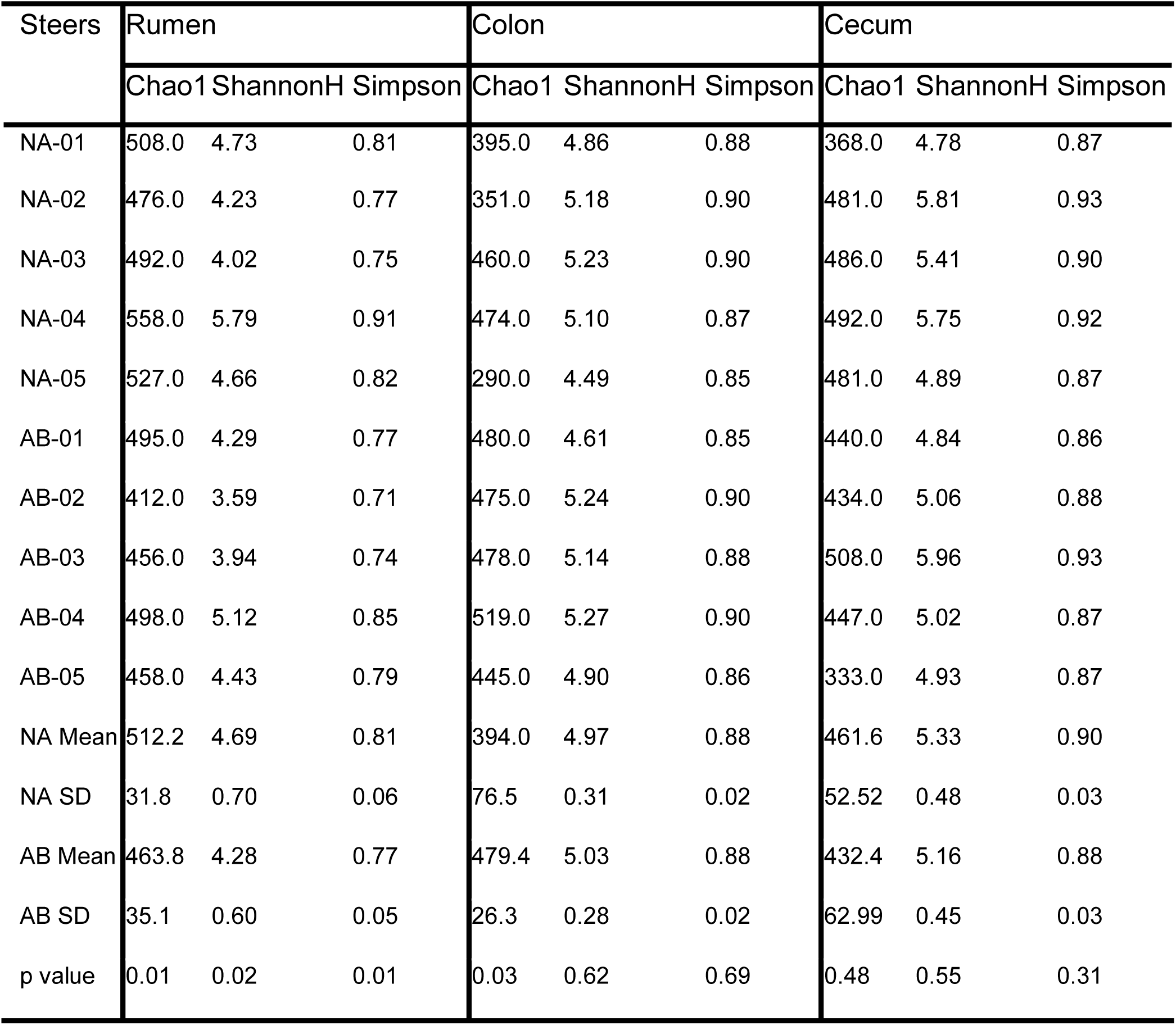
Alpha-diversity of the microbiome in the GI tract of feedlot cattle. Samples were collected from rumen, colon, and cecum of the untreated (NA) and treated (AB) steers. Untreated steers had a more diversified microbial population in the rumen.

### β diversity

The diversity among samples was calculated using Bray-Curtis index which measures the dissimilarity between the samples (Fig 1a). Although a consistent pattern is lacking for between-subject variation, few of the NA steers (NA-03, NA-04, NA-05 in the rumen and NA-02, NA-05 in the colon) exhibited slightly higher diversity when compared to other animals. Additionally, the principal component analysis (PCA) revealed that the rumen samples, except for NA-04, clustered together and were separated from cecum and colon samples (Fig 1b).

**Figure 1.**
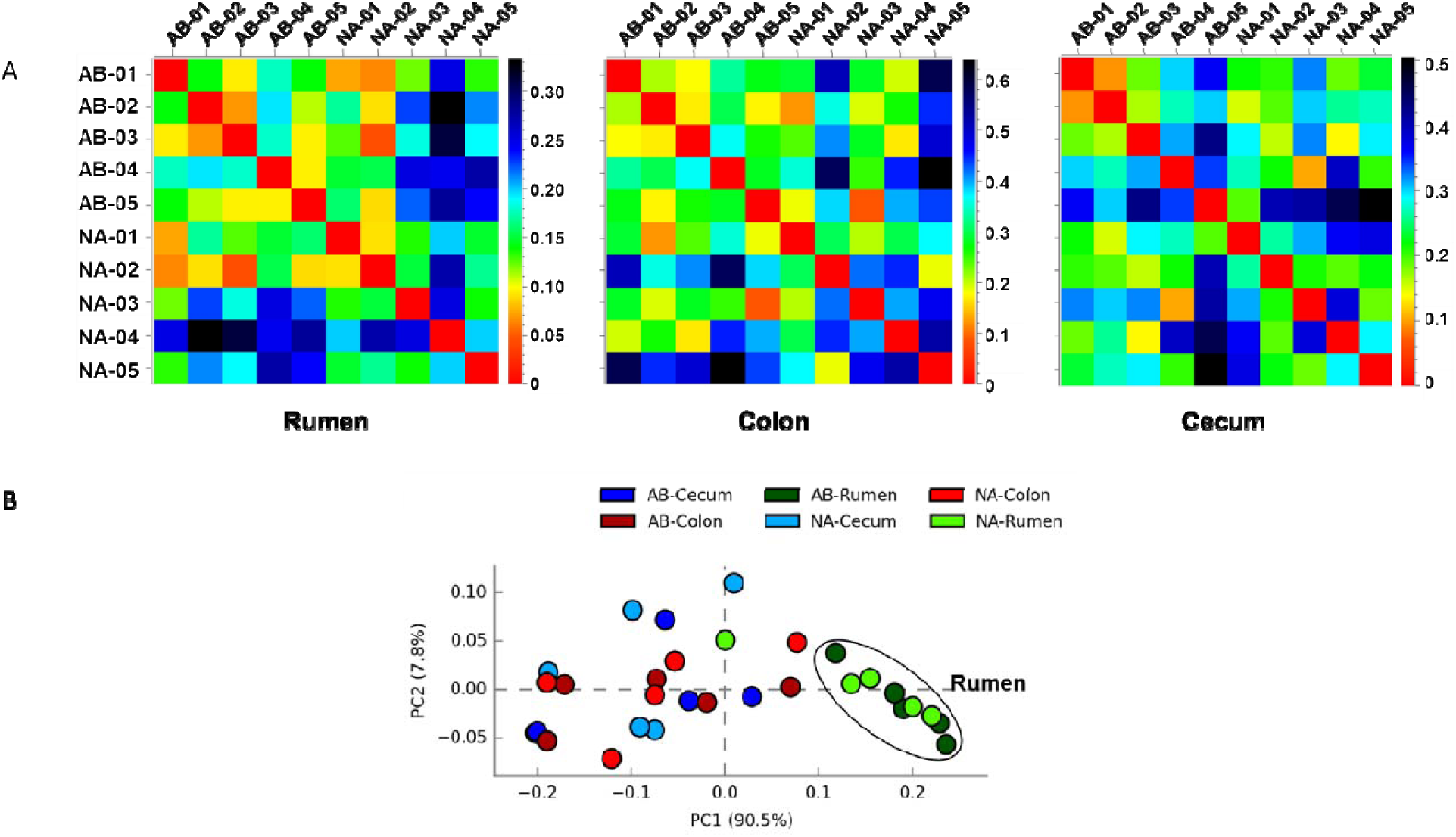
Microbiome diversity of bacterial population in rumen, colon, and cecum samples obtained from untreated (NA) and untreated (AB) steers. A) The beta-diversity of the bacterial community was measured using Bray-Curtis index. B) Principal coordinate analysis (PCA) of the microbial diversity in GI tract of steers showed that the rumen microbiome samples, except for NA-04, clustered together and separated from cecum and colon samples.

### Microbiome composition of the rumen, cecum, and colon at the phylum level

Samples from the rumen, colon, and cecum collected from NA and AB steers were analyzed for the phylum-level distribution of the bacterial population. In general, treatments had minimal effect on the distribution pattern of bacteria at the phylum level. However, the bacterial composition altered significantly between the locations in the GIT irrespective of the treatments. The percentage distribution of three major phyla - Bacteriodetes, Firmicutes, and Proteobacteria – are given in Fig 2. The most abundant phylum in rumen, cecum, and colon was Bacteriodetes. The abundance of Bacteriodetes in the rumen of AB steers was significantly higher (P < 0.05) compared to that in other compartments for both treatment groups but similar to that in the rumen of NA steers. The abundance of Firmicutes in the rumen of both NA and AB steers was lower (P < 0.05) than that in cecum and colon for both treatment groups. There were subtle changes in Bacteriodetes and Firmicutes distribution in the rumen of AB steers compared to that of NA steers, although these changes were not statistically significant. These alterations decreased the Firmicutes to Bacteriodetes ratio of 0.26 in NA steers to 0.20 in AB steers. There was no difference in the distribution of Proteobacteria between the treatment groups or between the three compartments of GIT.

**Figure 2.**
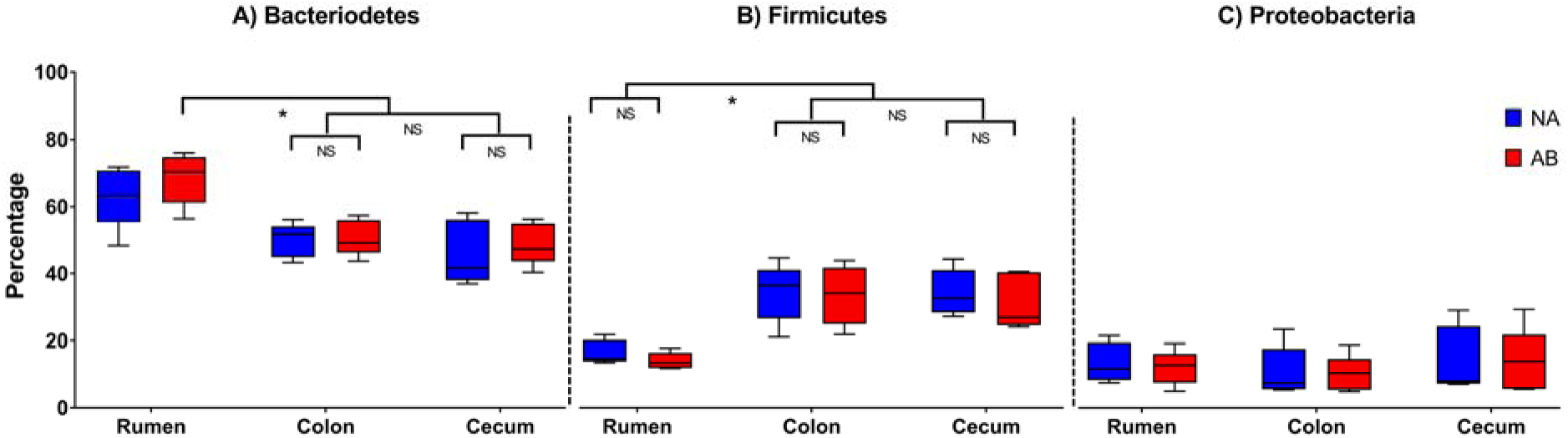
The phylum-level microbiome composition in the GI tract of steers. The treated (AB) steers received tylosin and monensin as feed additives while the untreated (NA) group received no antibiotic feed additives. There was no difference between the two groups within the GIT compartments for the three major phyla; A) Bacteriodetes, B) Firmicutes, and C) Proteobacteria. However, the percentage of Bacteriodetes and Firmicutes varied between the GIT compartments. In the rumen, the Firmicutes to Bacteriodetes ratio was lower for AB steers. The whiskers represent 10-90% confidence intervals and the solid black line represents the mean for the samples. NS denotes not significant; * denotes P < 0.05

### Microbiome composition of the rumen, cecum, and colon at the genus level

The shotgun metagenomics sequencing approach provides the potential for analyzing the bacterial community at a higher resolution. Even though the phylum-level distribution did not indicate major shifts in the bacterial community, in-depth analysis at genera-level suggested that treatment with antibiotic feed additives altered the microbial profile in the GIT of steers.

Distribution of most abundant genera in rumen, colon, and cecum are given in Fig 3. There were notable differences in the genus-level distribution between the GI tract compartments. Prevotella was the most abundant genus (39.82%) in the rumen while bacteriodes dominated in the cecum (23.89%) and colon (24.66%). Bacteriodes comprised only 18.46% of total population in the rumen. Genus Fibrobacter was markedly abundant in the rumen and contributed to 1.01% of rumen bacteria. In colon and cecum, Fibrobacter made up only 0.18% and 0.19% of the total population respectively. The other abundant genera in the rumen, but at a lower concentration in cecum and colon, were Paludibacter, Flavobacterium, Mitsuokela, Treponema, Pseudomonas, Selenomonas, and Bifidobacterium.

**Figure 3.**
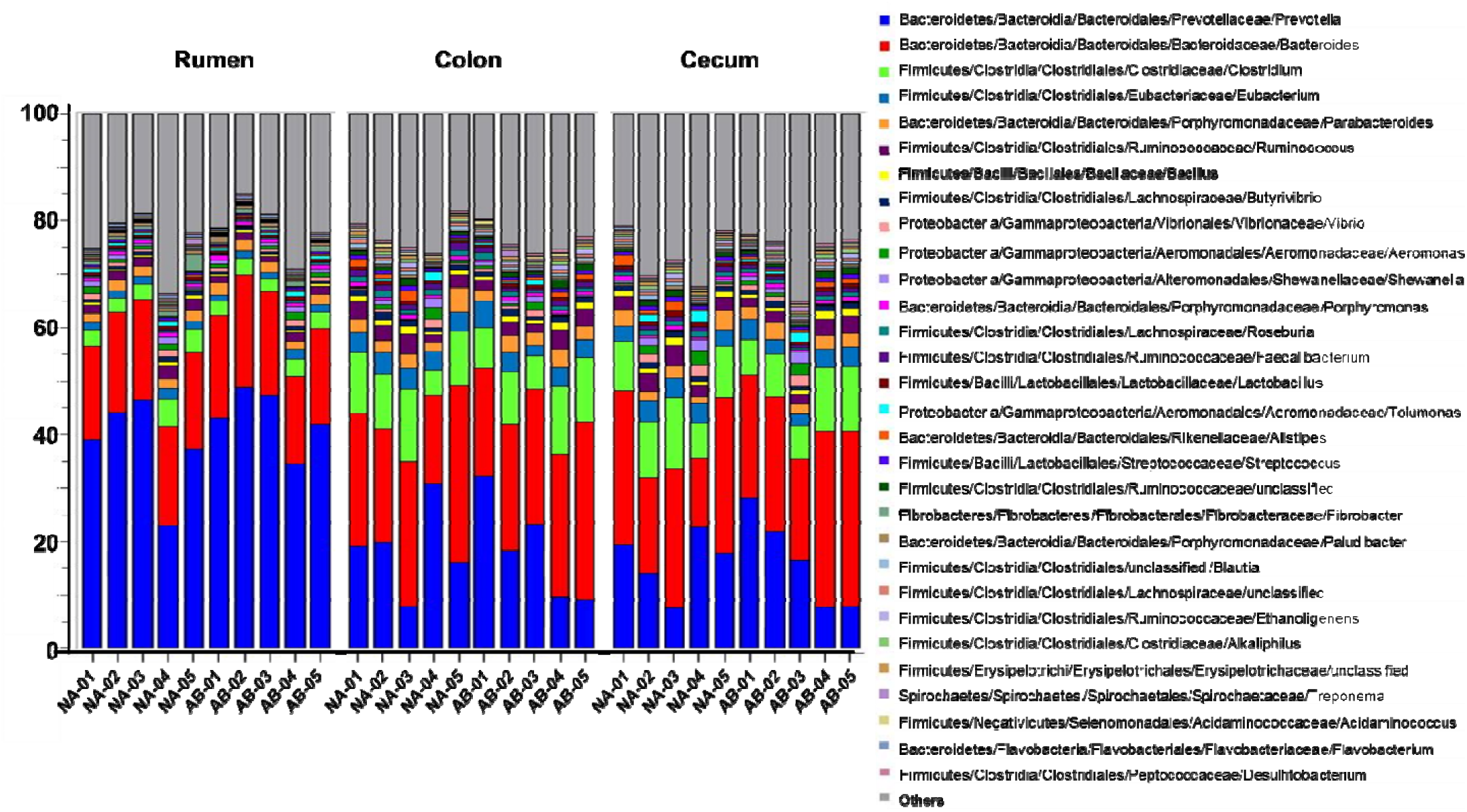
Genus-level distribution of the microbiome in GI tract of the steers. Phylogenetic distribution was generated in MG-RAST using RefSeq database with maximum e-value at 10^-5^ value, minimum percentage identity at 60 %, minimum alignment length of 30 amino acids and abundance 100. Each color represents one genus. In rumen samples, Prevotella was the most abundant genus, whereas, in colon and cecum, bacteriodes dominated.

To gain further insights, the statistical differences in genera that belong to Bacteriodetes and Firmicutes between NA and AB groups were analyzed using STAMP software. In the rumen, four genera belonging to Bacteriodetes – Spirosoma, Dyadobacter, Leadbetterella, and Zunongwangia – were significantly more abundant in the NA steers (Fig 4A). On the contrary, there was no difference in the distribution of Bacteriodetes in cecum and colon between treatment groups. Among the Firmicutes, few genera showed a significant difference between treatment groups in the rumen, colon, and cecum (Fig 4B, C, and D). In the rumen, gram-positive Firmicutes such as Ruminococcus and Fecalibacterium were reduced in the AB steers and were partially replaced by gram-negative members of Negativicutes. These changes were not surprising due to the selective action of monensin and tylosin against gram-positive bacteria. In the colon, gram positive Catenibacterium and unclassified Erysipelotrichacea and in the cecum, unclassified Lachnospiraceae were reduced significantly in AB steers. Similar to the rumen, members belonging to gram-negative Negativicutes increased in abundance in cecum and colon. Interestingly, gram-positive thermophilic organisms belonging to Thermoanaerobacteraceae family also increased in cecum and colon of AB steers. Additionally, in the cecum of AB steers, the abundance of gram-negative genera Cellulosilyticum and Syntrophomonas and thermophilic gram-positive genus Geobacillus also increased. The reason for the resistance pattern exhibited by gram-positive thermophilic organisms to tylosin and monensin is currently unknown.

**Figure 4.**
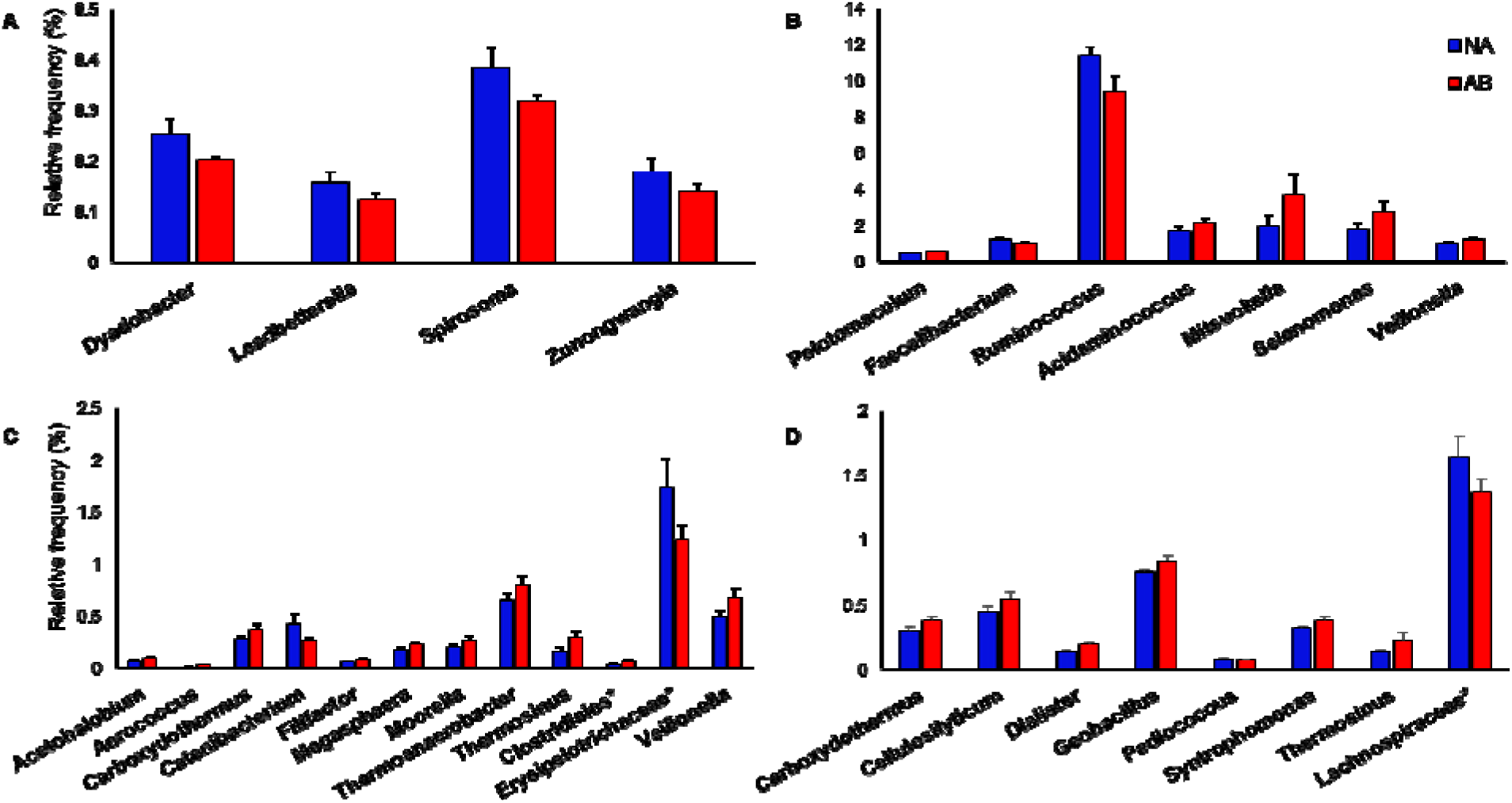
Alterations in abundance of the bacterial genera in the GI tract of feedlot steers following treatment. The statistical difference in genera distribution between untreated (NA) and treated (AB) steers was analyzed using STAMP software. Only those genera that are statistically different (p > 0.05) in abundance are given above. A) genera under Bacteriodetes in the rumen B) genera under Firmicutes in the rumen C) genera under Firmicutes in the colon and D) genera under Firmicutes in the cecum. The genera under Bacteriodetes were similar in both groups of animals in cecum and colon. * represents unclassified reads derived from families.

### Functional analysis of microbiome changes in the GI tract of feedlot cattle

The analysis of enzyme functions provides a metabolic blueprint of the bacterial community. The functional changes of the microbiome in the rumen, colon, and cecum of feedlot steers fed with and without antibiotic feed additives were predicted using SEED subsystems at level 2 hierarchy in MG-RAST pipeline (Fig 5). Overall, genes with unknown (null) functions were 24.17%, 23.81%, and 23.17% in the rumen, colon, and cecum respectively suggesting that functions of a large proportion of the genes that encodes for enzymes are yet unknown. Genes associated with protein biosynthesis were the most abundant among the known functions and the percentage was 7.91, 8.61 and 8.89 in the rumen, colon, and cecum respectively. The other dominant functions were associated with plant-prokaryote associations, RNA processing and modification, DNA repair, and central carbohydrate metabolism.

**Figure 5.**
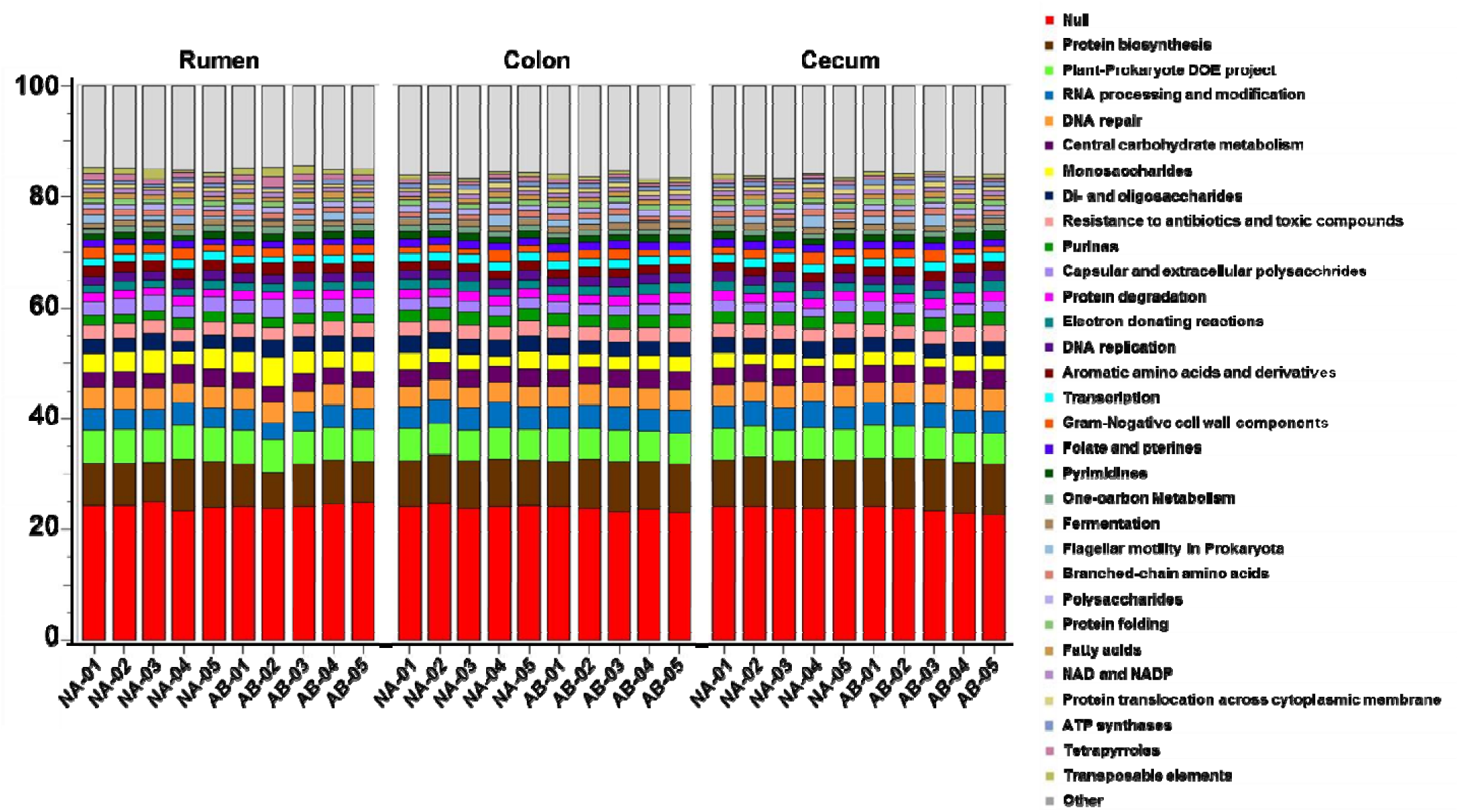
Predicted functional profile of the microbiome in the GI tract of the steers. The metagenomic functional analysis was performed in MG-RAST using Subsystems database with maximum e-value at 10^-5^ value, minimum percentage identity at 60 %, minimum alignment length of 30 amino acids and abundance 100. Treated steers (AB) were given antibiotic feed additives and untreated steers (NA) were the control animals which received no treatments.

The diversity in the functional characteristics of the metagenomes was analyzed using STAMP software. The PCA analysis of functional hierarchy showed clustering of all rumen samples, except NA-04, and segregating away from cecum and colon samples (Fig 6A). The cecum and colon samples did not have a definite pattern of segregation and were overlapping each other. This observation for functional analysis was similar to that found for taxonomical clustering at the genera-level.

**Figure 6.**
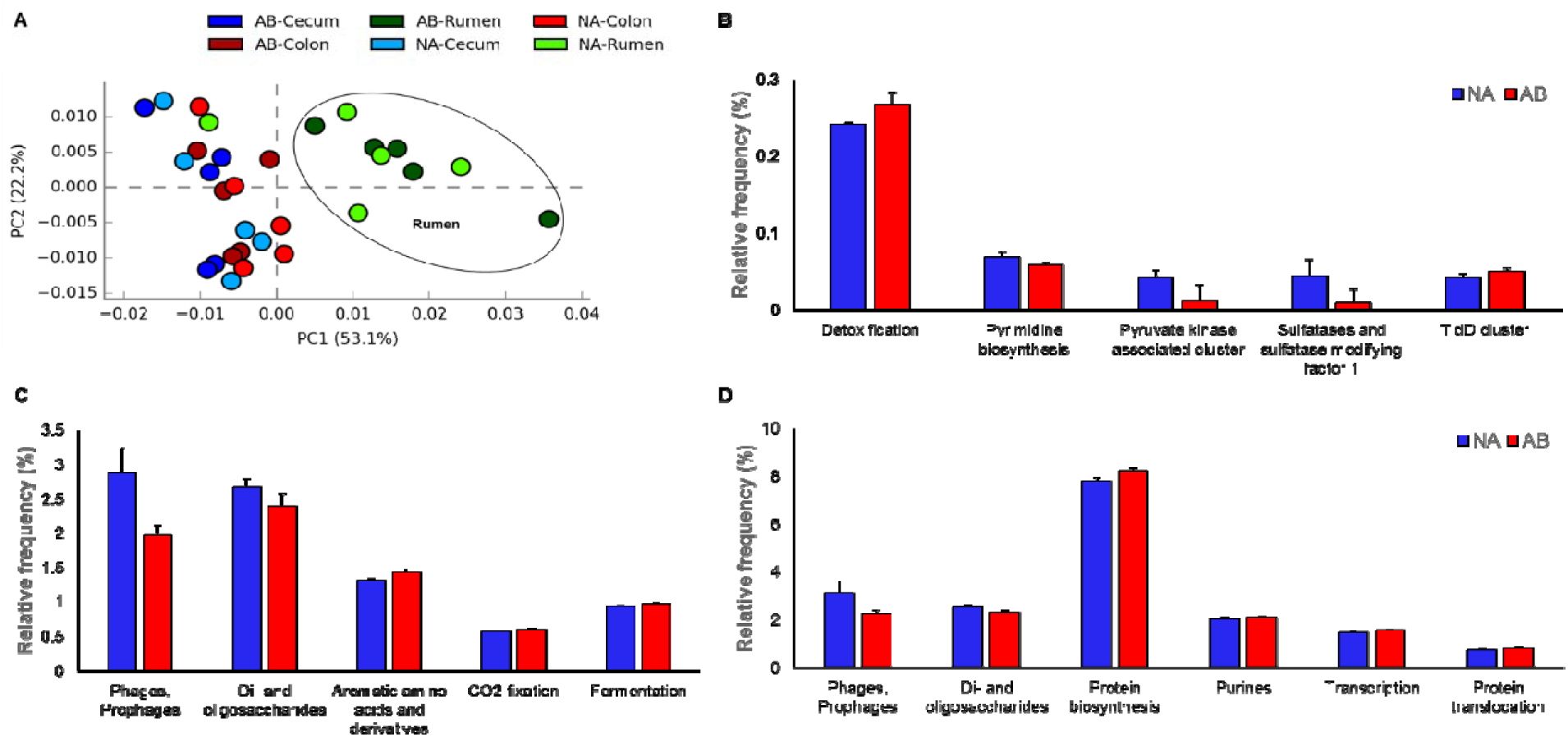
A) Principal coordinate analysis (PCA) of the functional diversity in the GI tract of steers. Treated steers were given antibiotic feed additives and untreated steers were the control animals. The rumen microbiome samples, except for NA-04, clustered together and separated from cecum and colon samples. Comparison of functional genes that are significantly different (p > 0.05) between NA (blue bars) and AB steers (red bars) B) in rumen C) colon and D) cecum are given above.

Further in-depth analysis revealed that the genes involved in detoxification were significantly increased in the rumen of AB steers (Fig 6B). However, sulfatases and sulfatase modifying factor 1 and pyruvate kinase associated genes were decreased in this group. In colon (Fig 6C) and cecum (Fig 6D), the genes associated to phages and prophages as well as di- and oligo-saccharides metabolism was reduced in the AB steers.

### Resistome composition of the gut

The presence of antibiotic resistance (AMR) genes in the pathogenic bacteria has been a threat to human and animal health. Microbiome of livestock could act as a potential reservoir for AMR genes for the pathogens. Antibiotic resistome of the GIT is composed of all the AMR genes present in pathogenic and non-pathogenic bacteria. The use of antibiotics as feed additives in livestock farming has been a controversial subject because of the possibility of gut microbiome gaining AMR determinants and expanding the antibiotic resistome. Functional metagenomics provides a potential resource for detecting the existence of AMR genes and antibiotic resistome composition in the gut microbial community.

In this study, the treated steers were provided tylosin and monensin. We compared the presence of AMR genes in the rumen, cecum, and colon of NA and AB steers (Fig 7) to determine whether antibiotic feed additives could influence the presence of AMR genes and expand the antibiotic resistome. The resistance genes pattern was predicted by BLAST searching assembled metagenome contigs against the ResFinder database. The rumen microbiota of three AB steers had the presence of the macrolide resistance genes *ermF* and *ermG*. However, these genes were not detected in the rumen of NA steers or in the cecum and colon of both groups. On the contrary, aminoglycoside resistance genes *aadE* and *aph(3)-III* were present only in the rumen of two NA steers, which were never provided antibiotics. Furthermore, Lincosamide resistance gene (*InuC*) was found in the rumen and colon of both treatment groups. Different variants of tetracycline resistance genes were also ubiquitously found in the rumen, cecum, and colon, irrespective of the treatment given during the feeding trial. Interestingly, the betalactam resistance genes were found only in the cecum and colon of both groups of steers but not in the rumen. Conversely, resistance gene *nimJ* against metronidazole was observed only in the rumen, irrespective of the treatments. The distribution of resistance genes in both group of steers points to the universal presence of antimicrobial resistance in the microbial population rather than resistance induced by treatments. Overall, no correlations were observed between the presence of AMR genes in the GIT microbiome and the administration of antibiotic feed additives.

**Figure 7.**
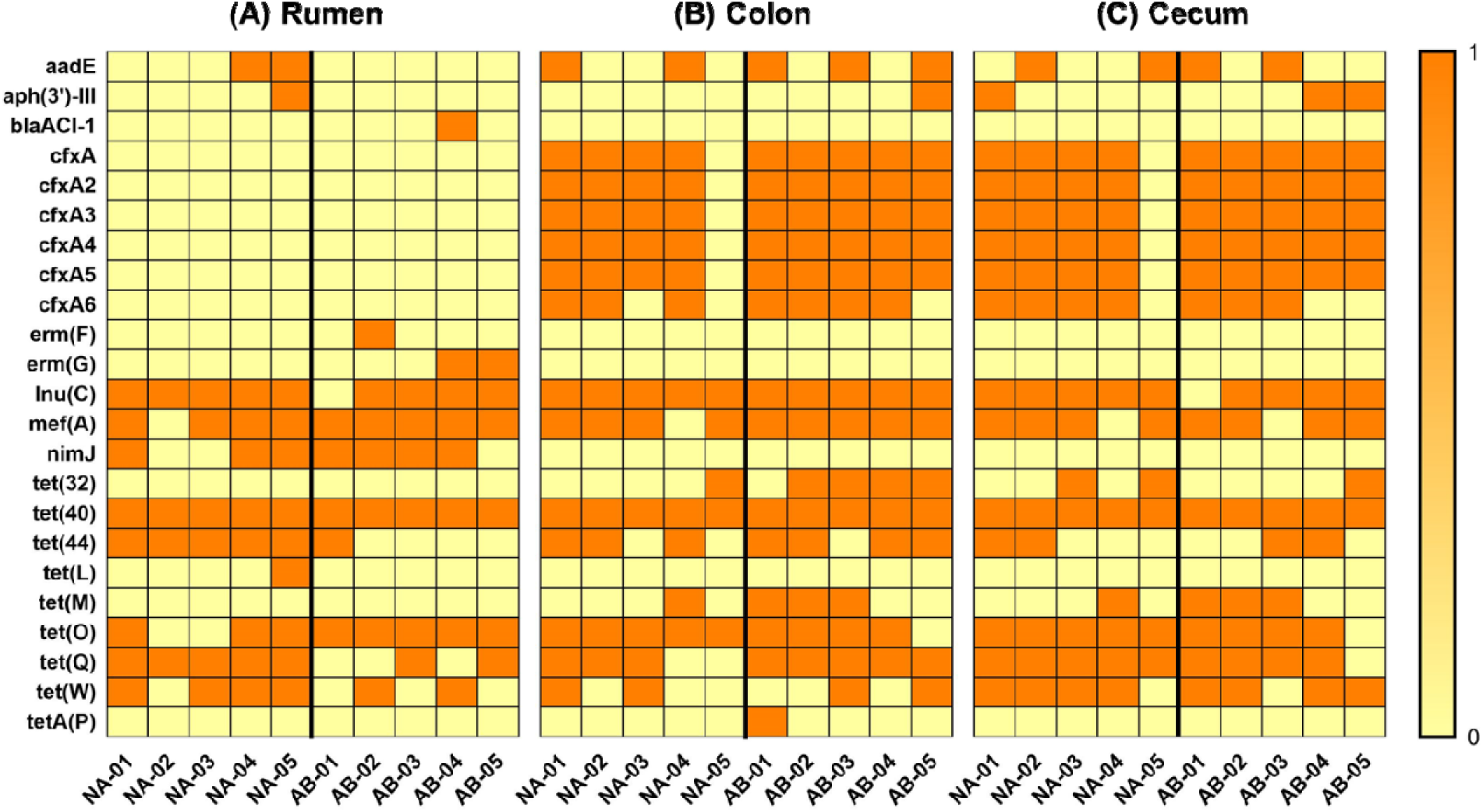
Heatmap showing the resistome profile in the GI tract of feedlot steers. Steers were fed Monensin and Tylosin as feed additives (AB) or control animals were not given any treatments (NA). Samples were collected from rumen, colon and cecum and DNA was subjected to metagenome sequencing using Illumina platform. The raw files were de novo assembled and antibiotic resistance genes were identified searching the assembled contigs against a local copy of ResFinder database using BLAST. The yellow color represent absence of the antibiotic resistance genes and orange color indicates presence. The macrolide resistance genes erm(F) and erm(G) were detected only in rumen of 3 AB steers.

## Discussion

This study compared the GIT microbiome and resistome of steers provided antibiotic feed additives, monensin and tylosin, with control animals that were fed the same ration but not provided antibiotics. The treated steers also received growth promoting implants and a beta-agonist during last 31 days of the trial, both of which are common practices in conventional feedlot management. Monensin, an anti-bacterial ionophore, is widely used in the feedlot industry to enhance production. It reduces the average feed intake and increases weight gain by enhancing propionic acid production in the rumen thereby resulting in improved energy efficiency ^31,32^. This feed additive also selectively reduces the gram-positive bacterial population. Tylosin is also an antimicrobial agent that acts against gram-positive bacteria. The major role of tylosin is to reduce the incidence of liver abscesses when steers are fed a rapidly fermentable carbohydrate-rich diet during the finishing phase of beef production. ^9–11^. When fed a high concentrate diet, the rumen pH is lowered due to the production of short-chain fatty acids, particularly lactate. Reduced pH promotes proliferation of *Fusobacterium necrophorum*, the primary agent that causes liver abscesses. Tylosin apparently inhibits the growth of *F. necrophorum* in the rumen and thus reduces the liver abscess incidences ^^7^,8^. Often monensin and tylosin are provided concurrently to cattle in North American feedlots. The active ingredient in Optaflexx is ractopamine-HCl and has been found to improve average daily gain, feed efficiency, and carcass gain ^33,34^. However, there is no available information that could link Optaflexx administration and the microbial pattern in the gut. The Revalor**^®^**-200 implants, which has anabolic steroids as active ingredients, is administered to feedlot cattle to improve their growth performance and some carcass traits ^35^. Recent findings suggest that both endogenous and exogenous estrogen and testosterone levels could influence the composition of gut microbiome ^36–38^. If growth promoting implants could influence gut microbiome, those changes could not be discerned from that caused by antibiotic feed additives. However, we speculate that antibiotics could produce a larger and more direct impact on the microbial composition compared to anabolic steroids. This is further supported by our finding that changes in the microbial dynamics in the rumen, cecum, and colon align with the activity of monensin and tylosin.

In the rumen, use of antibiotic feed additives was inversely associated with species diversity and richness as indicated by lower Chao1, ShannonH, and inverse Simpson indices for AB steers. Previous research also had shown that monensin decreased rumen bacterial diversity both *in vitro* and *in vivo* ^39,40^. Contrary to the findings in the rumen, the microbial population in colon and cecum samples had similar diversity for both treatment groups. This suggests that antibiotic feed additives had only a limited effect on the distal gut microbiota of steers. A primary reason could be that monensin is actively absorbed from the rumen, metabolized in the liver, and excreted through bile and only <10% of unmetabolized monensin reaches the distal gut ^41^. Similar to our findings, feeding monensin to lactating Holstein cows did not change the level of gram-positive and gram-negative associated 16S rRNA gene sequences in the colon-derived library ^23^.

The diversity indices and PCA profiles show that the microbial community in the rumen is different from that of distal gut irrespective of the use of antimicrobial feed additives. Studies conducted in dairy cows and a Nellore steer also found that the microbial population was significantly different between forestomach, small intestine, and large intestine ^21,22^. Results from our study further corroborate their finding that GIT region is a strong determinant of microbial community structure and also that microbial population in the fecal samples may not be representative of forestomach microbiota. The composition of the microbial population in the rumen is significantly influenced by the diet ^15,16,42^. In forage fed cattle, the community is more diverse and tend to have a higher Firmicutes to Bacteriodetes ratio. Major phyla that dominate in the rumen of forage-fed animals include Firmicutes, Bacteriodetes, Fibrobacter, and Proteobacteria. As cattle transition from high forage to high concentrated diets, the diversity is lost and the composition altered ^42,43^. However, changes in the microbial community due to high grain diet do not follow a definite pattern. Few studies have reported that the abundance of Firmicutes increased in the rumen of dairy cows when fed with a highly fermentable diet that induced acidosis ^6,43^. On the other hand, the abundance of Bacteriodetes in the rumen increased in beef steers when 60% or more of the diet was composed of grain ^16^. In the present study, > 90% of the diet was derived from corn and corn byproducts that were highly fermentable. The percentage of reads belonging to phylum Bacteriodetes in the rumen of NA and AB steers were 63.07 ± 9.19 and 68.47 ± 7.72 respectively. Furthermore, the Firmicutes constituted only 16.56 ± 3.53 % and 14.04 ± 2.49 % of the reads in NA and AB steers. Additionally, the Bacteriodetes predominated in both treatment groups in the cecum and colon but the percentage was lower when compared to the rumen. Our findings suggest that a high concentrate diet provides a conducive environment for the survival of the Bacteriodetes. Although there were no significant differences in the read count percentage between the two treatments, AB steers had lower Firmicutes to Bacteriodetes ratio (0.26 vs 0.20) in the rumen. This shift in the ratio is in accordance with the activity of monensin and tylosin, which inhibits the growth of gram-positive Firmicutes. The ratio observed in this study was lower than those observed in other dietary studies, which analyzed the microbial pattern in the rumen ^16,43^. This could be due to the difference in the quantity of concentrate fed to the cattle in these experiments. Previous experiments used a lower proportion of concentrates (< 80%) in the diet and hence the shift in microbial dynamics could be less pronounced than in the present study.

The human microbiome of a healthy adult has a higher proportion of Firmicutes than Bacteriodetes ^44^. An increased Firmicutes to Bacteriodetes ratio is observed in overweight and obese people and is associated with a Western diet, which has more rapidly digestible carbohydrate content ^45^. Conversely, a decrease in the ratio is correlated with weight loss ^46,47^. However, these findings in human may not be applicable in ruminants because of the anatomical and physiological differences. Ruminants depend on gluconeogenesis for up to 90% of the total glucose requirements of the body ^48^. Propionic acid, a major precursor for glucose, is actively absorbed from the rumen and is converted to glucose in the liver. Phylum Bacteriodetes, in general, produces acetate and propionate as end-products of fermentation ^49^. Therefore, a higher proportion of Bacteriodetes in the rumen could be favorable for efficient feed utilization and body weight gain.

At the genera-level, Prevotella was the most common genus in the rumen and second-most abundant genus in cecum and colon. This was in accordance with several studies which described Prevotella as the most abundant Bacteriodetes member found in rumen ^5,15,16,22,25,50^. It is also one of the core members found in rumen microbial population ^5^,^50^ irrespective of diet and location. Furthermore, the proportion of Prevotella in the rumen could increase up to 10 times in grain-fed cattle when compared to forage-fed animals ^5,16,18,51^. Prevotella could utilize lactic acid and convert it into propionic acid. Thus, an increase in abundance of Prevotella is beneficial for high concentrate-fed cattle. Additionally, it was found that the abundance of Megasphaera and Selenomonas populations also increase when cattle are gradually adapted to a high concentrate diet ^16,18,43^. Megasphaera and Selenomonas are members of Veilonellacea family and are propionic acid producers. In this study, the use of antimicrobial feed additives further increased the abundance of Megasphaera, Selenomonas, Mitsuokella, and Veilonella in the rumen. Even in cecum and colon, unclassified members of <u>Erysipelotrichaceae</u> and Lachnospiraceae were reduced and partially replaced by members of <u>Veillonellaceae</u> family. This shift in microbial population could prevent the accumulation of lactic acid and also increase the propionic acid synthesis.

Another core rumen microbial population is the genus Ruminococcus, which is composed of cellulolytic and lactic acid producers ^5^. Previous research has shown that the abundance of Ruminococcus decreased when animals were transferred from a forage-based diet to a high grain diet ^5,6,18^. We observed that monensin and tylosin further reduced the abundance of Ruminococcus genus in the rumen. This will be beneficial for cattle that are maintained on a high concentrate diet because of the decreased lactic acid production and associated risk of acidosis. However, the reduction in abundance of genera Ruminococcus, <u>Erysipelotrichaceae</u> and Lachanospira in various compartments of AB steers may impact the ability of GIT pathogen exclusion capacity of these animals. Various studies in humans have reported that these genera are important in suppressing the growth of enteric pathogens in the gut^3,52^.

We also characterized the gene functions of the bacteria using level 2 Hierarchical classification in MG-RAST. Our findings indicated that there are functional differences between the GIT regions. Similar variability in gene functions was observed in the GIT of the dairy cattle ^22^. The most abundant gene functions were related to protein synthesis, RNA processing, DNA repair, carbohydrate metabolism, and resistance to antimicrobials and toxic compounds. These functions are essential for the survival and proliferation of the microbial community. When the antibiotic feed additives were included in the diet, steers had a higher abundance of genes associated with detoxification in the rumen. However, genes associated with detoxification formed less than 0.3% of the total functional genes and could be of lesser impact. In cecum and colon, the use of monensin and tylosin is correlated with reduced copy of genes for di- and oligosaccharide metabolism and genes associated with phage.

In this study, we also characterized the antimicrobial resistance (AMR) genes present in the GIT of the feedlot animals. Recent experiments have used shotgun metagenome sequencing as a diagnostic tool to predict AMR in swine ^53^, environment ^54,55^, bovine ^56,57^, and human ^58^ associated microbiome. Environmental samples collected from beef pens and abattoir revealed AMR determinants against 17 classes of drugs ^54^. Similarly, fecal samples obtained from 6 beef heifers possessed AMR determinants against multiple drugs ^56^. This study would be the first to report AMR determinants in the GIT of feedlot animals. There were AMR determinants against 8 classes of drugs in the various compartments. One of the major findings was that use of tylosin did not increase the presence of AMR genes in the AB steers. Tylosin resistance genes *tlr, ermB*, and *ermX* were not detected in any of the samples. Although macrolide resistance genes *ermF* and *ermG* were present only in the rumen of AB steers, the gene encoding for macrolide efflux pump, *mefA*, was ubiquitously found in all three compartments of both treatment groups. This is in contrast to the earlier finding where the use of tylosin increased the presence of macrolide resistance genes in feedlot steers^59,60^. Another finding was that the distribution of *cfxA* and *nimJ* genes, which provide resistance to beta-lactams and metronidazole respectively, differed between forestomach and hind-gut. This could indicate that sampling feces alone might not reveal the whole picture on AMR in the case of ruminants. Sampling forestomach along with fecal samples would be more effective in predicting a comprehensive AMR profile although collecting rumen samples would be difficult in farm conditions. Overall, our results did not show significant difference in AMR profile of steers in NA and AB groups. Since the total duration of antibiotic feed additive in this study was 74 days, the AMR profile we discovered here may not be representation of longer term effect of such feed additives.

## Materials and Methods

### Ethics Statement

All procedures and protocols used in this study were reviewed and approved by the Institutional Animal Care and Use Committee (IACUC) at the South Dakota State University, Brookings, SD.

### Animal experiments

A subset of 10 Simmental x Angus crossbred steers raised from a common herd were selected for analysis. Five steers were sampled per treatment, which included; non-antibiotic treated cattle (NA) and implanted cattle fed a beta-agonist and antibiotic feed additives (AB). Cattle were housed and managed according to the approved Institutional Animal Care and Use Committee protocol. Prior to weaning, steer calves were managed at the SDSU Antelope Field Station near Buffalo, S.D. where they were randomly designated to treatment after being stratified by cow age, calf birth date, and weight. Steers within the AB group received calf hood implants (36 mg zeranol; Ralgro, Merck Animal Health) at an average 74 days of age. After weaning, all steers received the same diet during morning feed delivery. Steers were backgrounded for a total of 71 days on a high roughage ration (grass hay, concentrate pellets, dry corn cobs, glycerin, distillers grains, limestone, and minerals). During backgrounding, treated steers received a Revalor**^®^**-IS implant (80 mg trenbolone acetate and 16 mg estradiol; Merck Animal Health).

### Feedlot Management

To be finished, steers were maintained within assigned feedlot pens and acclimated to the finishing diet using five step-up diets over a period of 68 days. The final finishing diet was composed of 40% wet corn gluten feed (Sweet Bran, Cargill Inc., Blair, NE), 48% dry-rolled corn, 7% grass hay, and 5% supplement containing 58.25% ground corn, 29.57% limestone, 5.59% iodized salt, 4.65% ammonium chloride, 0.93% trace mineral mix, 0.25% thiamine, and 0.21% Vitamins A, D, and E. Nutrient composition was 72.58% DM, 13.9% CP, 2.04 Mcal/kg NEm, and 1.38 Mcal/kg NEg. Steers in the AB group received 478.3 g/ton monensin (Monensin 90, Elanco Animal Health, Greenfield, IN) and 96.1 g/ton tylosin (Tylosin 40, Elanco Animal Health), while steers in the NA group did not receive hormone implants, ionophores, beta-agonists, feed-grade or injectable antibiotics or antimicrobials.

During the final phase of the finishing period, steers were placed into a GrowSafe^®^ system. Steers within AB group were implanted with Revalor^®^-200 (200 mg trenbolone acetate/20 mg estradiol; Merck Animal Health). Additionally, steers in this group received ractopamine hydrochloride (Optaflexx^®^, Elanco Animal Health) during the last 31 days of finishing. All cattle had ad libitum access to fresh water throughout the study. At harvest, which was based on a target back fat thickness of 1.5 cm, the average age of NA steers were 13 months and AB steers were 14 months. The average dry matter intake (DMI) was greater for AB (12.58 kg ± 0.23) than NA (11.54 kg ± 0.24; *P* < 0.0001). Average daily gains (ADG) were also greater for AB (1.79 kg ± 0.05) than NA (1.54 ± 0.05; *P* < 0.0001).

### Sample collection and genomic DNA isolation

Samples of digesta were collected from rumen, colon, and cecum immediately after slaughter and were snap-frozen in liquid nitrogen. These samples were then transferred to −80^0^C and stored until use. Genomic DNA was isolated from these samples using Powersoil DNA isolation kit (Mo Bio Laboratories Inc, CA). Approximately 100 mg of sample contents were transferred to bead tubes for DNA extraction. After adding 60μl of solution C1 to the bead tubes, samples were homogenized for 2 min using Tissuelyser (Qiagen, MD). Remaining steps of DNA isolation was performed according to the manufacturer’s protocol. The DNA was eluted in 50 μL nuclease free water. The quality of isolated DNA was analyzed using NanoDrop™ One (Thermo Scientific™, DE) and DNA was stored at −20^0^C until use.

### Microbial DNA enrichment and sequencing

Selective enrichment of microbial genomic DNA was performed using NEBNext^®^ Microbiome DNA Enrichment Kit (New England Biolabs, Inc. MA) ^26^. The enrichment process was performed according to manufacturer’s protocol. Briefly, 0.5 μg of genomic DNA was treated with 80 μl of MBD2-Fcbound magnetic beads in the presence of binding buffer and incubated at room temperature for 15 min with rotation. After incubation, beads were separated by keeping the tubes on a magnetic rack for 5 min. The supernatant, which contained microbial DNA, was transferred to another tube and were purified using Agencourt AMPure XP beads (Beckman Coulter) and stored at −20^0^C.

Enriched microbial genomic DNA from test samples was used for shotgun metagenome sequencing. The concentrations of genomic DNA samples were measured using Qubit Fluorometer 3.0 (Invitrogen, Carlsbad, CA) and the concentration was adjusted to 0.2 ng/μl. After normalization, sequencing libraries were prepared using Nextera XT DNA Sample Prep Kit (Illumina Inc. San Diego, CA). Tagmentation of samples using 1 ng of template was conducted according to manufacturer’s protocol, followed by PCR amplification using a unique combination of barcode primers. PCR products were then cleaned up using Agencourt AMpure XP beads (Beckman Coulter). Purified products were normalized using library normalization protocol suggested by manufacturer. Equal volumes of normalized libraries were pooled together and diluted in hybridization buffer. The pooled libraries were heat denatured and spiked with 5% of the Illumina PhiX control DNA prior to loading the sequencer. Illumina paired-end sequencing was performed on the Miseq platform using a 2 x 250 paired-end sequencing chemistry.

### Sequence data processing

The raw data files were de-multiplexed and converted to FASTQ files using Casava v.1.8.2. (Illumina, Inc, San Diego, CA, USA). The raw sequence data for all the samples in this study has been deposited in NCBI Sequence Read Archive (NCBI SRA) under the bioproject number - PRJNA390551. FASTQ files were concatenated and analyzed using MG-RAST^61^. The sequences were subjected to quality control in MG-RAST. This included de-replication, removal of host specific sequences, ambiguous base filtering, and length filtering. The taxonomical abundance of organisms was analyzed using MG-RAST with Best Hit Classification approach using Refseq database and parameters were limited to minimum e-value of 10^−5^, minimum percentage identity of 60%, a minimum abundance of 100, and a minimum alignment length of 30 amino acids. The functional abundance was analyzed using Hierarchical Classification in MG-RAST using Subsystems and parameters were limited to minimum e-value of 10^−5^, minimum percentage identity of 60%, a minimum abundance of 100, and a minimum alignment length of 30. The OTU abundance tables were downloaded from MG-RAST and were used for downstream statistical analysis.

To determine the resistome composition of the samples, raw data files were assembled in CLC workbench 9.4 using *de novo* metagenome assembler (Qiagen Bioinformatics, CA). The assembled metagenome was searched against a local copy of ResFinder database^62^ using BLAST.

### Statistical analysis

Influence of AB and NA on animal performance (DMI and ADG) were evaluated using PROC MIXED in SAS. Cow age was used as a covariate and LS-means were computed for response variables. The significance level was set at P < 0.05. The phyla distribution was statistically analyzed using ANOVA with Tukey’s multiple comparison test. Indices of taxonomical diversity in the samples was determined using Explicet software^63^. Bootstrapping was performed to reduce the biases in sequencing depth. A total of 50 bootstraps were conducted to calculate the α- diversity indices –Shannon, Simpson, and Chao1. Diversity between samples (β- diversity) was estimated by deriving Bray-Curtis index. Statistical differences among Firmicutes and Bacteriodetes in various digestive tract segments were calculated using STAMP software ^64^. The differences were estimated using two-sided Welch’s t-test. Similar parameters were used for analyzing the statistical differences among overall functional abundance. Corrected p-values below 0.05 were considered significant. The principal component analysis (PCA) of gene functions and taxonomical distribution were also generated using STAMP software.

## Acknowledgements

This work was supported in part by the USDA National Institute of Food and Agriculture, Hatch projects SD00H532-14 and SD00R540-15, and South Dakota Beef Industry Council grant awarded to JS. The funding agencies had no role in study design, data collection, and interpretation, or the decision to submit the work for publication.

## Author Contributions

JS, MT, AB, and KO conceived and designed the experiments. MT, MW, SG, GJF and ATF performed the experiments. MT analyzed the data. JS, AB, KO, DB and JCH contributed reagents and materials. MT and JS wrote the manuscript. All authors reviewed and approved the manuscript.

